# Microbial Thermal Response Strategies Impact Environmental Fitness of Horizontally Transmitted Symbiont Strains

**DOI:** 10.64898/2026.02.19.706762

**Authors:** Alexander Nuckols, Patrick T. Stillson, Alison Ravenscraft, Nicole M. Gerardo

**Affiliations:** Department of Biology, Emory University, Atlanta, GA, USA; Department of Biology, University of Texas at Arlington, Arlington, TX, USA

**Keywords:** *Caballeronia*, Gene expression, Heat shock, Heteroptera, Symbiosis, Thermal stress

## Abstract

Microbial symbionts in insect-microbe mutualisms are critical for host survival and fitness. However, symbioses are sensitive to external stressors. Understanding how a warming climate will impact these associations, therefore, is critical. Previous work on horizontally transmitted bug-*Caballeronia* bacterial symbionts has shown that different symbiont strains confer variable host outcomes under thermal stress, suggesting that environmental symbiont acquisition could be advantageous under shifting temperature regimes. However, it is unknown what specific bacterial thermal responses could impact microbial survival in the environment, which would be key to host acquisition. We evaluated *in vitro* thermal response mechanisms of heat-vulnerable and heat-resistant strains of *Caballeronia* using microbial transcripts, to identify pathways that may impact the symbionts’ respective thermal optima. We found that a heat-resistant strain prioritizes induction of thermally stable outer membrane components, motility structures, and molecular chaperones, allowing it to increase growth through induction of central metabolism and protein synthesis. Meanwhile, a heat-vulnerable strain arrests growth, favoring induction of genes related to alternative metabolism and biofilm formation. These results are indicative of variable bacterial thermal response strategies for elevated temperature survival. Ultimately these responses may alter which symbionts are more available to insect hosts as temperatures continue to rise.

**Importance:** Many eukaryotes host beneficial microbes that synthesize essential nutrients or can break down harmful compounds. While important, if these microbes are acquired from the local environment (*i.e.,* soils or plant surfaces), external stressors can destabilize the associations by impacting microbial growth, limiting the pool of available microbes. Using the bug-*Caballeronia* model system, we analyzed the *in vitro* thermal stress response of two *Caballeronia* symbiont strains that have different thermal optima. We found that stress responses involving increased motility, thermally stable membrane synthesis, and molecular chaperones may increase growth under thermal stress in symbionts. These results indicate that different thermal stress coping strategies may be favored in bug-*Caballeronia* system under increasing global temperatures. More broadly, these results provide insight into how microbial stress responses could shape adaptation of symbioses to a warming climate.

## Introduction

Many eukaryotes engage in mutualistic associations with microbes. These microbial symbionts are often essential to the fitness of their hosts, aiding in nutrient breakdown and synthesis, chemical detoxification, and defense against parasites and pathogens (1–4). However, external stressors such as high temperatures can reduce the benefits that symbionts confer to their hosts (5–7), with some stressors transforming mutualistic relationships into antagonistic (8).

To persist under the various external stressors that both endosymbiotic and environmental microbes are exposed to, bacteria have adapted various stress responses. In the case of increased temperatures, bacteria commonly have heat shock pathways that maintain protein stability and cellular function with a global regulatory response, activating chaperone proteins that aid in protein-folding at high temperatures, and proteases to break down and recycle denatured proteins (9). High temperatures have also been associated with diverse bacterial transcriptional responses, including alterations in expression of genes related to biosynthesis and metabolic pathways, motility, and numerous other cellular processes (10,11). In addition to these variable regulatory responses, closely related species of bacteria can have different thermal optima. These differential thermal optima are the result of evolutionary trade-offs in thermal tolerance, which can ensure community survival, especially in the case of fluctuating environmental conditions and temperatures (12).

For microbes, living in the environment requires numerous stress responses that could influence evolutionary trade-offs of different symbiont transmission modes of animal hosts. Vertically transmitted symbionts are passed directly from parent to offspring, which can result in a dramatic reduction of the symbiont genome. With little external influence, these symbionts rapidly acquire mutations in genes that are unnecessary for symbiosis or survival within a host. Stress response genes are often among those lost in this genomic reduction (13,14). While the host is usually able to provide a stable, relatively stress-free environment, vertically transmitted symbionts are susceptible to elevated stress, like high temperatures, resulting in host failure when the symbiont is eliminated (15). Horizontal symbiont transmission occurs when a symbiont is acquired anew each generation, either from conspecific hosts in the population, or from the environment directly. Unlike vertically transmitted symbionts, environmentally acquired, free-living microbes must retain adaptive genes necessary to survive under local conditions and to compete with other microbes (16). The retention of genes necessary for a free-living lifestyle suggests that environmentally derived symbionts likely have a suite of stress response mechanisms, offering a wide range of potential adaptive benefits to hosts under varying climatic conditions (17–19). Additionally, environmental acquisition allows intergenerational symbiont exchange, allowing for rapid acquisition of novel symbionts that are better adapted to the current environmental conditions. This suggests that environmental acquisition may provide greater benefits for hosts, especially in the face of rapidly changing climates.

True bugs (Hemiptera) in the heteropteran superfamilies Coreoidea, Lygaeoidea, and Pyrrhocoroidea (20,21) acquire their microbial symbiont *Caballeronia* anew each generation from their environment during the second instar (developmental stage) (22,23). *Caballeronia* live in pouches in the host midgut called crypts, and provide essential amino acids and B vitamins to the host (24). When reared without *Caballeronia*, hosts have high mortality rates, smaller body size, and delayed development (23,25). Previous work has identified that a number of genes are involved in establishing and maintaining this symbiosis. Key genes include those whose products influence motility, stress responses, cell wall synthesis, o-antigen biosynthesis (26).

By promoting an increase in host weight and faster development, or by conferring insecticide resistance when acquiring insecticide-degrading *Caballeronia* strains (3,27), symbiont identity can impact host fitness (28). Symbiont identity can also impact host outcomes under different temperatures. Eastern leaf-footed bugs (*Leptoglossus phyllopus*), for example, were inoculated with six different *Caballeronia* strains with varying thermal optima, then reared from 24–36°C. While symbiont *in vitro* thermal optima did not reliably predict host outcomes under thermal stress, differences in host mortality, average body size, and development rate were observed between hosts with different symbionts. These differences became more extreme at high temperatures, with one strain, which was sensitive to high temperature *in vitro*, entirely failing the host at 36°C (19). This suggests that, if symbionts are adapted to local temperatures, environmental symbiont acquisition could provide an advantage to the hosts under thermal stress.

To investigate mechanisms by which different symbionts can impact host outcomes under thermal stress, two of the six strains used previously were selected based on differences in their *in vitro* thermal optima and variation in host conferred benefits: V-LZ003 (heat-vulnerable; lower thermal optimum: 24°C, failed host at high temperatures) and R-LZ019 (heat-resistant; higher thermal optimum: 28°C, benefitted host at high temperatures) (19). These symbionts show differential host outcomes under thermal stress, aligning with their *in vitro* thermal optima. At 24°C, the two strains did not differ in host outcomes, however at 36°C, insects inoculated with the vulnerable strain suffered increased mortality and reduced body size compared to hosts with the resistant strain. In a previous study, *In vivo* (*i.e.,* within host) transcripts of these two symbiont strains were analyzed across different temperatures, with each symbiont strain expressing different *in vivo* transcriptional profiles across temperatures. Both strains downregulated significantly more genes at high temperatures than they upregulated, however, the vulnerable symbiont differentially expressed 31% fewer genes than the resistant symbiont between 24°C and 36°C. Among those upregulated by both strains were protein-folding genes. Meanwhile, the resistant strain showed significant upregulation of heat shock and flagellar genes not observed in the vulnerable strain (29). This *in vivo* transcriptomic analysis identified key genes that may contribute to the observed variation in host outcomes at higher temperatures when provided with different symbiont strains.

While host outcomes are important, symbiont fitness in the environment is also critical for environmentally acquired symbioses. Increasing temperature regimes should select for microbes in the environment with greater ability to protect themselves from thermal stress. Through investigating *in vitro* thermal response mechanisms, we attempt to elucidate how different symbiont responses to elevated temperatures may affect the survival of symbiotic microbes in the environment, and subsequently how this change might affect host acquisition of symbionts. To uncover the different thermal response strategies of V-LZ003 and R-LZ019, we performed a transcriptomic analysis to identify each strain’s transcriptional response when cultured at 24°C or 36°C in liquid media. V-LZ003 and R-LZ019 have thermal optima below 36°C, so culturing at this temperature puts both strains under some degree of thermal stress. Our results reveal vastly different thermal response strategies, indicating that higher thermal tolerance is associated with the induction of metabolic processes, molecular chaperones, and synthesis of thermally stable membrane components, as opposed to alternative metabolism and community behaviors, such as biofilm formation.

## Materials and Methods

### Bacterial culturing

Two *Caballeronia* species V-LZ003 (heat-vulnerable) and R-LZ019 (heat-resistant) were used due to their different *in vitro* thermal optima and host insect outcomes under thermal stress (19) (Accessions: V-LZ003: GCF_031451465.1; R-LZ019: GCF_031450825.1). Liquid cultures were grown overnight in YG (yeast-glucose) broth at 28°C, then diluted 1:20 in fresh YG broth (30). The diluted cultures were then grown at 24°C, 30°C, and 36°C until they reached an optical density (OD) of 0.6-0.7. Cultures were then divided into 1 mL aliquots, pelleted, and the broth was removed. Five hundred µL of RNAlater (Invitrogen, Waltham, MA, USA) was added, and cells were stored at -20°C before RNA extraction.

### RNA extraction and sequencing

Total RNA from the liquid cultures was extracted for each sample (n = 3 per strain/treatment) using the Promega SV total RNA isolation system protocols for Gram-negative bacteria (Madison, WI, USA). Total RNA was sequenced by Novogene (Beijing, China), if it passed a minimum RNA integrity number ≥ 6.0, using their Prokaryotic RNA library preparation pipeline, which removed bacterial rRNA. Sequencing was performed on a NovaSeq PE150 (Illumina, San Diego, CA, USA) with approximately 2 Gb of raw reads produced for each sample.

### Mapping reads

We trimmed the raw reads with Cutadapt v.5.1 to remove both adapter sequences and low-quality sequences. Reads with low quality (quality cutoff set to 20), a minimum length < 20 bp, or N > 10% were dropped from all analyses (31). We mapped the trimmed and filtered reads to each strain’s respective genome using HISAT2 v.2.2.1 (32) and counted mapped reads for each gene using featureCounts v.2.1.1 with the default settings (33).

### Differential Expression

Differential expression analysis was performed using DESeq2 for each strain in each of the three temperature treatment groups. The 30°C samples were found to overlap with other temperature treatments based on the principal component analysis and had minimal gene count variation when compared to either the 24°C or 36°C treatments. Therefore, the 30°C samples were removed from subsequent analyses. For comparisons between symbiont species, only differential expression for orthologous genes, present in both strains, was analyzed. For comparisons within species, each species’ entire respective genome was used for analysis.

### Gene Ontology and KEGG Enrichment Analyses

Using the R package “GO_MWU” (https://github.com/z0on/GO_MWU), we calculated the log2(foldchange) * -log(adjusted P value) for all transcripts in gene ontology (GO) enrichment analyses to consider both magnitude and significance of differential expression for each GO term (34). Genes from each species’ genome were matched with their respective KEGG terms using eggNOG Mapper v.2.1.8 (35,36) followed with a KEGG enrichment analysis (function = ‘enrichKEGG’, package = ‘clusterProfiler’) (37). Significant KEGG pathways were visualized, using log2(fold change) for each significant mapped transcript (P value < 0.05) (package = ‘pathview’) (38).

### Motility assay

Eight cultures each of V-LZ003 and R-LZ019 were grown overnight in YG broth at 24°C and 36°C with shaking. When cultures were in exponential phase, we measured their OD_600_ with a spectrophotometer, then centrifuged and resuspended them in YG broth to an OD_600_ of 3.0. Two µL of these cell suspensions were stabbed into the center of 0.3% agar YG plates. Two replicates were conducted for each culture (8 cultures x 2 replicates each). The plates were sealed with Parafilm to prevent dehydration and incubated right side up for 15 h at 24°C or 36°C, corresponding with the temperatures at which each culture was initially grown. The diameter at the widest point of the turbidity zone was measured with a ruler. Because the vulnerable symbiont was nonmotile, we used a two-sample t-test to evaluate differences between distance traveled for the resistant symbiont under each temperature condition.

### Biofilm formation assay

Eight cultures each of V-LZ003 and R-LZ019 were grown overnight in YG broth at 24°C and 36°C with shaking. When cultures were in exponential phase, we measured their OD_600_ with a spectrophotometer and diluted them in YG broth to an OD_600_ of 0.01. One hundred µL of each dilution was added to the wells of a 96-well microtiter plate. Two replicates were used for each culture (8 cultures x 2 replicates). Cultures raised at 24°C were added to one microtiter plate and those raised at 36°C were added to another. Microtiter plates were incubated without shaking for 48 h at 24°C and 36°C, corresponding with the temperatures at which each culture was initially grown.

After 48 h, cultures were dumped out, and the microtiter plates were rinsed with water. One hundred and twenty-five µL of 0.1% crystal violet solution was added to each well. The plates were incubated for 15 min at room temperature before the crystal violet solution was disposed, and the plates were again rinsed with water. Microtiter plates were left to dry at room temperature overnight. Then, 125 µL of 33% acetic acid was added to each well, and the plates were incubated for 15 min at room temperature. A plate reader was used to measure the absorbance of each well at 550 nm, with 33% acetic acid used as a blank. The blank absorbance was subtracted from each sample to determine the OD_550_. We used a Kruskal-Wallis test followed by pairwise Dunn’s test to determine significance between OD_550_ for the symbionts under each temperature condition. For all R based analyses, v4.5.1 was used (39).

## Results

### Transcript statistics

Reference transcriptomes were constructed for both symbionts by mapping the transcripts against the corresponding symbiont genome for all 18 samples. There was an average of 22.4 (SD = 3.1, range = 9.1) million reads mapped for V-LZ003 and an average of 26.5 (SD = 2.8, range = 9.6) million reads mapped for R-LZ019 (Table S1).

### Transcriptional variation across temperatures

Before evaluating differences in expression between the two symbionts, we performed a principal component analysis to look at expression variation across temperature treatments. This confirmed that different temperature treatments primarily clustered among themselves, indicating that transcriptional profiles in each strain shift as temperatures increase. Heat-resistant strain R-LZ019 had tight clustering for both 24°C and 36°C treatments, with the 30°C samples containing one outlier (Fig 1A), while heat-vulnerable strain, V-LZ003, had the tightest clustering among the 36°C samples, with both the 24°C and 30°C samples each containing one outlier (Fig 1B). Due to the significant overlap with other temperature treatments of the 30°C samples for both strains, the 30°C treatment group was excluded from subsequent analyses.

**Figure 1.**
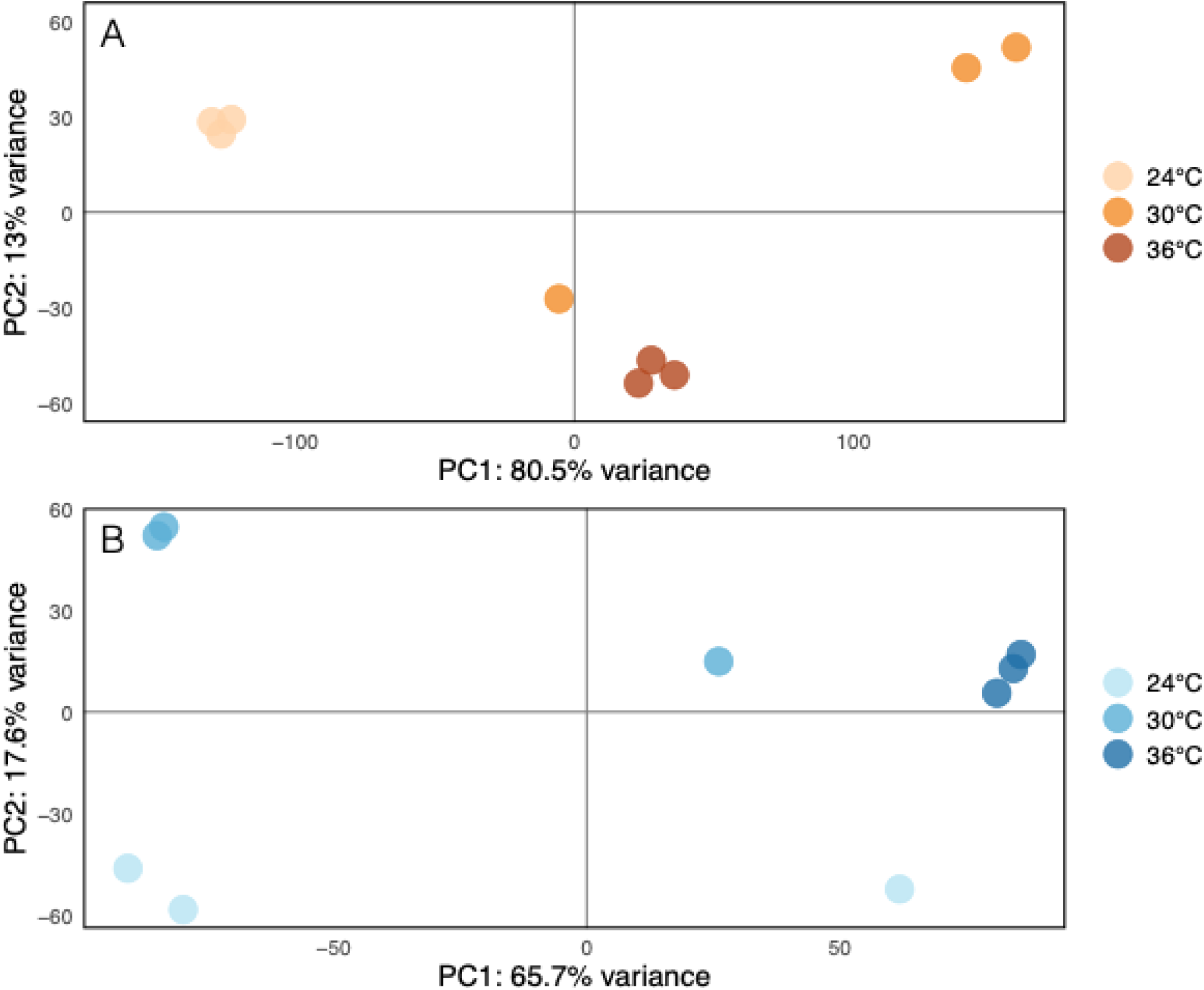
Principal component analysis based on gene expression reveals intersample variation at the intermediate temperature. A principal component analysis was performed on the DESeq2 output and plotted to visualize sample variation across treatment groups for (A) the resistant strain, R-LZ019, and (B) the vulnerable strain, V-LZ003. In both strains, the 30°C treatment groups had high sample variation.

To assess how each strain responds to thermal stress, we compared the transcriptomes of the two *Caballeronia* symbionts under the 24 and 36°C temperature treatments. Differential expression analysis (FDR corrected P value < 0.05 with a log2(fold change) ≥ |1|) revealed that the heat-resistant strain upregulated 48% more genes than it downregulated at 36°C compared to 24°C, while the heat-vulnerable strain downregulated 98% more genes than were upregulated at 36°C compared to 24°C (Table 1). Across temperature treatments for the resistant symbiont, the fold change was much larger for the upregulated genes than downregulated genes, indicating more extreme induction of genes in response to high temperatures (Fig 2A), while the vulnerable strain had an increase in repression under thermal stress conditions (Fig 2B). Transcriptional differences between strains for each temperature treatment were analyzed with differential expression analysis using a filtered symbiont genome containing conserved genes between the two (3868 genes). For both temperature treatments, a similar number of genes were differentially expressed in each direction and with similar fold changes (Table 1; Fig 2C, D).

**Figure 2.**
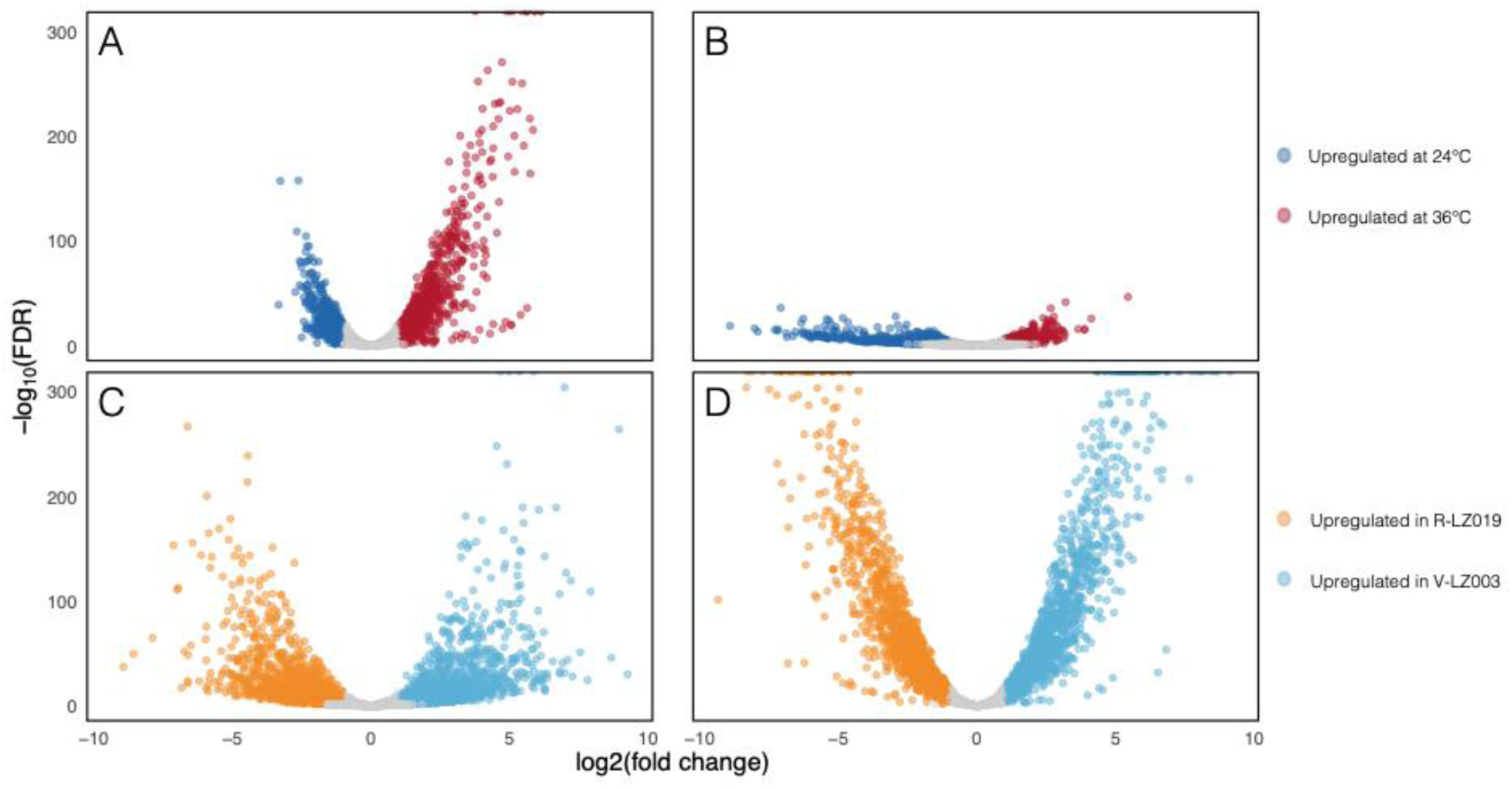
Heat-resistant and heat-vulnerable strains have different global transcriptional responses to increased temperature. Symbiont transcriptional variation depicted with volcano plots for (A) 24°C–36°C thermal comparison in the resistant strain, (B) 24°C–36°C thermal comparison in the vulnerable strain, (C) vulnerable–resistant symbiont comparison at 24°C for orthologous genes, and (D) vulnerable–resistant symbiont comparison at 36°C for orthologous genes. Colored points indicate differentially expressed transcripts by pairwise comparison (FDR-corrected P value < 0.05 with a log2(fold change) ≥ |1|).

**Table 1.**
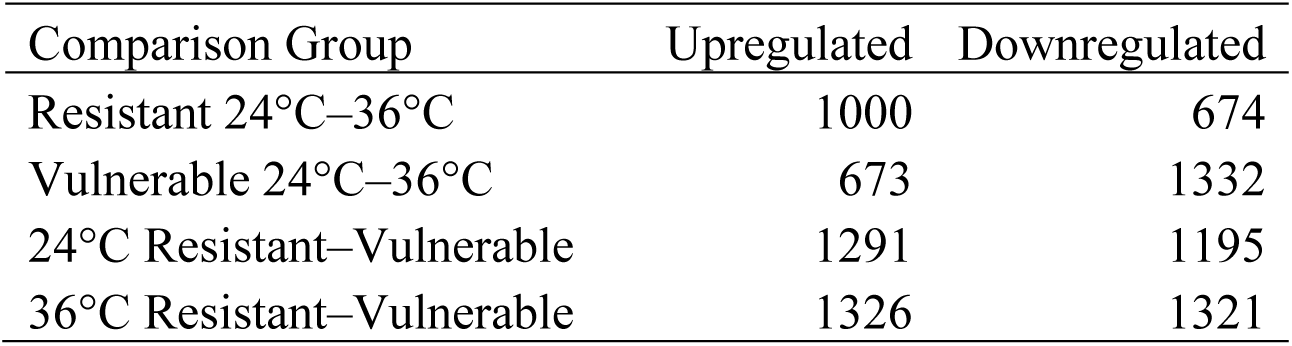
Number of differentially expressed genes between each of the four comparison groups. The direction of differential expression reflects expression at 36°C compared to 24°C for rows 1 and 2, and expression in the heat-vulnerable strain compared with the heat-resistant strain for rows 3 and 4.

### Identification of genes and pathways associated with symbiont thermal responses *in vitro*

A rank-based gene ontology enrichment analysis was performed to identify differential patterns of functional enrichment between the two strains with varying responses across temperature treatments. The heat-resistant strain upregulated numerous metabolic and biosynthetic processes at 36°C compared to 24°C (Fig 3A). ATP metabolism, amide biosynthesis, and ribonucleoside triphosphate biosynthesis were among the highest upregulated categories, indicating induction of energy metabolism, transcription, and translation. Other highly upregulated GO terms include amino acid activation, cell motility, and locomotion. The few processes downregulated include those for transport pathways, signal transduction, and regulation of metabolism.

**Figure 3.**
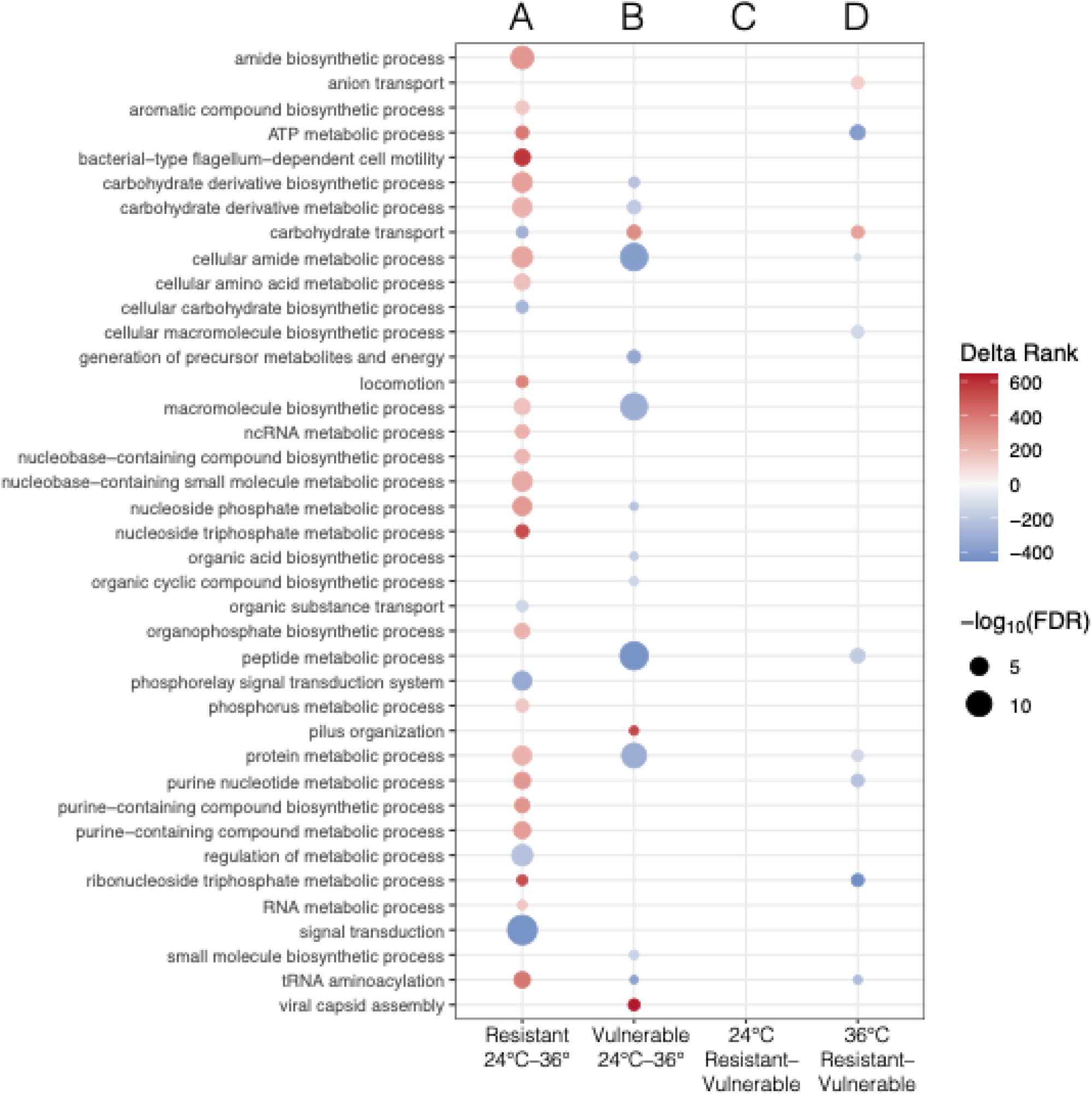
Resistant strain induces and vulnerable strain represses biological processes at high temperature. Dot plot showing delta rank and significance of enriched GO terms among each comparison group (P value < 0.01). Delta rank indicates the degree of up-/down- regulation for each GO term. Red indicates upregulation and blue indicates downregulation. Resistant strain, R-LZ019, mostly upregulates biological processes at the higher temperature (A, red circles). The vulnerable strain, V-LZ003, mostly downregulates these processes at higher temperatures (B, blue circles). For the 24°C comparison between strains (C), no significant GO terms were identified.

The heat-vulnerable strain had half as many significantly differentially expressed GO terms (Fig 3B). In contrast with the resistant strain, many biosynthetic and metabolic processes were downregulated at 36°C. The most highly downregulated processes at elevated temperatures were cellular amide and peptide metabolisms, suggesting a decrease in protein synthesis at higher temperatures. Additionally, generation of precursor metabolites and energy and tRNA aminoacylation were downregulated. Only carbohydrate transport, pilus assembly, and viral capsid assembly were found to be upregulated at the higher temperature.

When comparing orthologous gene expression between strains, we found a moderate, but significant, negative correlation (P value < 0.001), indicating that orthologs generally have opposite regulation patterns in response to thermal stress (Fig S1). No biological processes were significantly differentially expressed at 24°C between strains (Fig 3C), while 10 biological processes were differentially expressed at 36°C, consistent with observed patterns found within strains (Fig 3D). Broadly, biosynthetic and metabolic processes had lower expression in the vulnerable strain at 36°C, while transport process related genes increased expression.

Finally, KEGG enrichment analysis revealed several significant pathways that were differentially expressed within strains across temperature treatments. Most pathways had heterogeneous regulation patterns, with some genes being induced and some repressed. To determine general direction and strength of regulation, we calculated the median log2(fold change) of all genes in each pathway. A total of 46 significant divergent pathways were identified (P value < 0.05) (Table S2). Divergent pathways were determined based on each strain’s median log2(fold change) among genes in that pathway. Strains that had opposite median log2(fold changes) were considered divergent. Pathways with large observed differences in median log2(fold change) between strains were of particular interest. These include biosynthesis of nucleotide sugars, central metabolic pathways, flagellar assembly, oxidative phosphorylation, and ribosomal components, all of which were upregulated in the resistant strain and downregulated in the vulnerable strain.

### Motility assay

The GO and KEGG enrichments pointed to upregulation of motility in the resistant strain. Thus, we evaluated this with a swimming motility assay. The vulnerable strain showed no motility in any replicates at either temperature. Meanwhile, the resistant strain was motile at both temperatures but traveled 37% farther at 36°C (P value < 0.001) (Fig 4).

**Figure 4.**
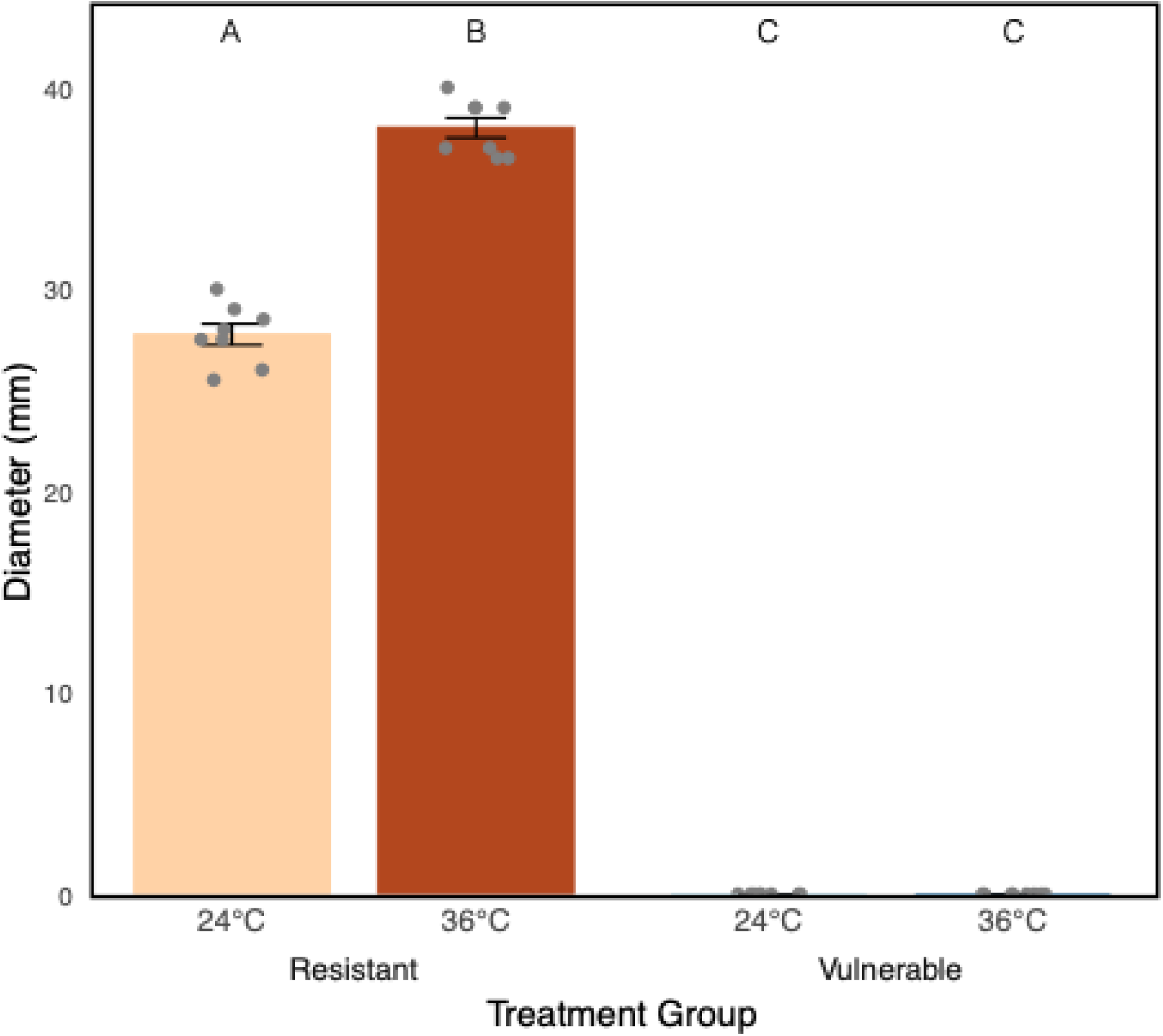
Vulnerable strain is non-motile. Bars indicate the average diameter (mm) of the turbidity zone (± SEM) on 0.3% agar YG plates after 15 h for each treatment group, representing the bacteria’s swimming motility. Points represent biological replicates (n = 8). Different letters above bars indicate significant differences between groups.

### Biofilm formation assay

The KEGG enrichment also indicated that the vulnerable strain upregulates key components of the biofilm formation pathway, so we evaluated this with a biofilm formation assay. We found higher biofilm formation in the vulnerable strain compared to the resistant strain at both temperatures, with 474% greater biofilm formation at 24°C (P value < 0.01) and 592% greater biofilm formation at 36°C (P value < 0.001). Both strains showed similar biofilm formation between temperature treatments (V-LZ003 P value = 1.00; R-LZ019 P value = 1.00) (Fig 5).

**Figure 5.**
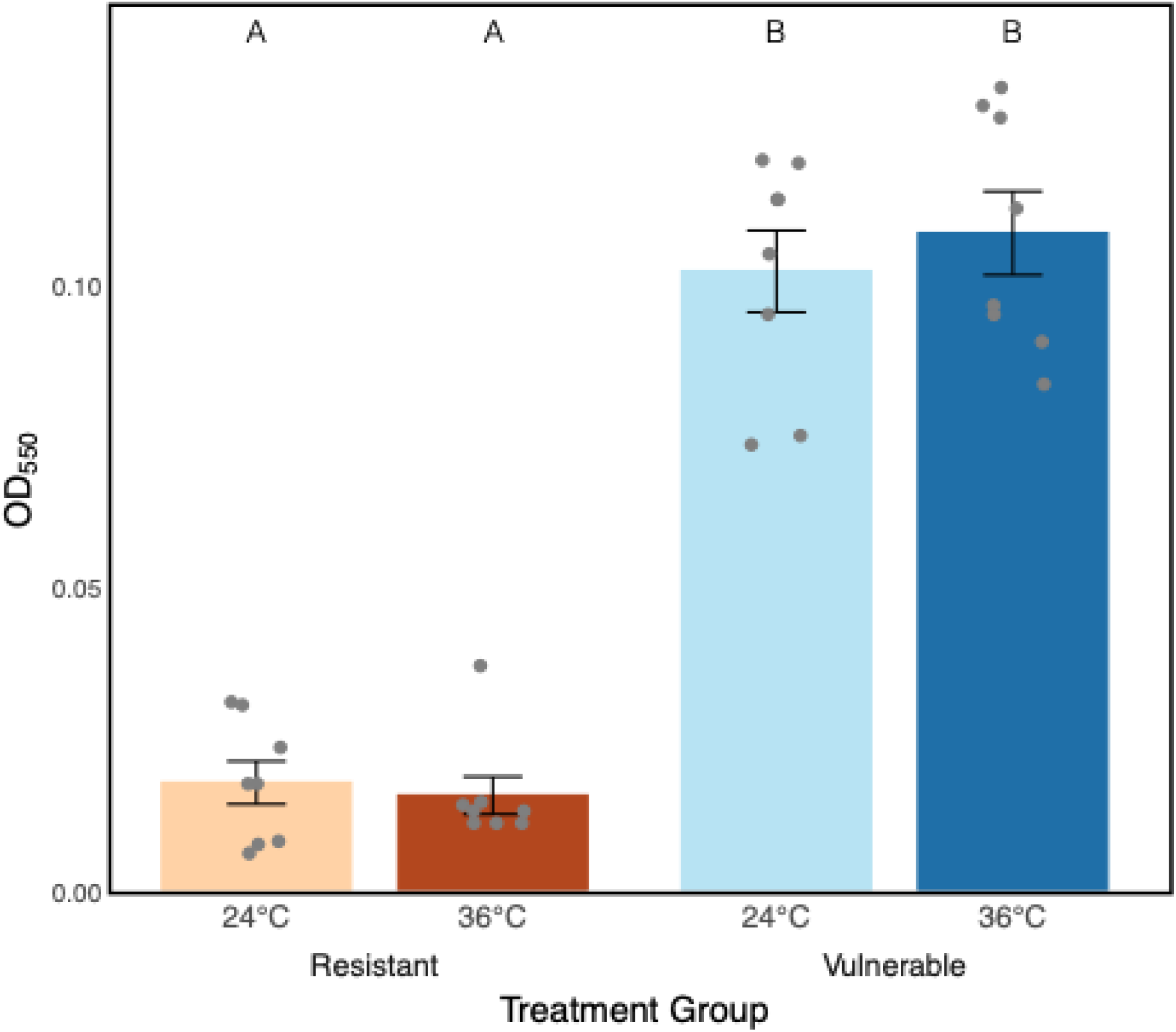
Vulnerable strain has higher biofilm formation than resistant strain. Bars indicate the average OD_550_ (± SEM) of cultures stained with crystal violet after 48 h growth, representing the bacteria’s total biofilm formation. Points represent biological replicates (n = 8). Different letters above bars indicate significant differences between groups.

## Discussion

Through an analysis of *in vitro* transcripts for two different symbiont strains with vastly different *in vivo* thermal stress responses, we found that the thermally resistant strain had higher gene induction at 36°C compared to 24°C, while the thermally vulnerable strain had higher repression, particularly for genes involved in biosynthetic and metabolic processes. These results were similar to *in vivo* findings (29) and consistent with our hypothesis and with the resistant strain’s higher thermal optimum. Several studies on *E. coli* have associated transcriptional upregulation of biosynthetic and metabolic processes with higher growth rates, especially for genes involved in nucleotide and protein synthesis (40–42). This is consistent with our GO enrichment analysis, which identified amide and ribonucleoside triphosphate biosynthesis among the most upregulated processes in the resistant strain, and cellular amide and peptide biosynthesis as among the most downregulated processes in the vulnerable strain.

### Differential stress responses between strains

#### Cell proliferation

Transcriptional regulation patterns for several key pathways suggest that the vulnerable strain enters growth arrest under thermal stress conditions (36°C), while the resistant strain increases cell growth and proliferation at the same temperature. Firstly, translation machinery is upregulated in the resistant strain and downregulated in the vulnerable strain. KEGG pathway analysis revealed divergent regulation of ribosomal proteins between strains. At 36°C, the resistant strain highly upregulated all ribosomal small 30S subunit proteins and nearly all large 50S subunit proteins, while the vulnerable strain highly downregulated these same components. Repression of translation in the vulnerable strain was also achieved with a reduction in tRNA aminoacylation through induction of the *VapC* toxin, which has been shown to arrest bacterial growth through cleavage of tRNAs (43,44). The vulnerable strain upregulates *VapC*, arresting protein synthesis, while the resistant strain downregulates both *VapC* and *VapB*, the antitoxin that inhibits *VapC*. Induction of this toxin-antitoxin system contributes to tolerance of several types of stress (i.e. antibiotic, oxidative, and thermal) by arresting growth and promoting long-term survival (45–47). This suggests that the vulnerable strain is arresting growth through translational repression, potentially to improve its chances of long-term survival, while the resistant strain instead increases cell growth and proliferation. This variation between strains is further supported by the highly divergent regulation of cell division protein *FtsZ*, which is upregulated in the resistant strain and downregulated in the vulnerable strain (Table S3).

#### Energy Production

The resistant and vulnerable strains also exhibited large differences in their regulation of energy metabolism and respiration pathways when coping with thermal stress. At 36°C, key genes in all three central metabolic pathways—glycolysis, the pentose phosphate pathway, and the TCA cycle—are universally upregulated in the resistant strain and downregulated in the vulnerable strain. Median log2(fold change) values reflect this (Table S2). This pattern is consistent with the vulnerable strain’s apparent growth arrest and the resistant strain’s increase in growth at 36°C. The standard oxidative phosphorylation pathway is also upregulated in the resistant strain.

Component structures for ATP synthase, cytochrome c oxidase, cytochrome o ubiquinol oxidase, NADH dehydrogenase, and succinate dehydrogenase are upregulated in the resistant strain and downregulated in the vulnerable strain. Transcriptional differences in cytochrome o ubiquinol oxidase and NADH dehydrogenase are particularly extreme in both strains. Cytochrome o ubiquinol oxidase subunits I and III and NADH dehydrogenase subunits *M*, *L*, *J*, *NuoK*, *NuoI*, *NuoG*, and *NuoN* are among the most divergently regulated orthologs in the transcriptomes (Table S3). Meanwhile, the cytochrome bd complex is upregulated in the vulnerable strain and downregulated in the resistant strain. Within the cytochrome c reductase molecule, the cytochrome b and iron-sulfur subunits are downregulated in the resistant strain but upregulated in the vulnerable strain. However, the cytochrome c1 subunit is upregulated in the resistant strain and downregulated in the vulnerable strain (Fig S2). These results further demonstrate that the resistant strain upregulates standard metabolic pathways, with high induction of oxidative phosphorylation structures.

The vulnerable strain’s regulatory pattern suggests that it utilizes alternative electron donors for energy production. The vulnerable strain downregulated NADH dehydrogenase and succinate dehydrogenase. Bypassing early respiratory structures such as NADH dehydrogenase is common of alternate electron donors that use downstream structures as electron transport chain entry points.

When being utilized as alternative electron donors, sulfur compounds can be used to bypass NADH dehydrogenase and succinate dehydrogenase, and formate bypasses NADH dehydrogenase (48,49). The vulnerable strain’s upregulation of cytochrome c reductase components suggests this as a possible entry point for alternative electron donors into the electron transport chain. The KEGG sulfur metabolism pathway is upregulated in the vulnerable strain (Table S2). Specifically, genes involved in the breakdown of methionine, a sulfur-containing amino acid that would be present in the rich media, is upregulated. Additionally, genes involved in transporting extracellular sulfur compounds, such as the *CysFUWA* sulfate uptake complex and the *SsuACB* sulfonate transport complex, are upregulated, suggesting this as a possible alternative energy-generating mechanism induced in the vulnerable strain under thermal stress; however, genes that have been previously identified for use in bacterial sulfur oxidation, such as sulfide-quinone reductase, are not present in the V-LZ003 genome. The vulnerable strain also upregulates a number of other genes that are related to exploitation of alternative electron donors in other bacteria (50–53), including two 2-hydroxyacid dehydrogenase genes, seven acyl-CoA dehydrogenase genes, six aldehyde dehydrogenase genes, and seven xanthine dehydrogenase genes.

#### Thermal Stress Response Genes

Use of alternative electron donors has been associated with growth arrest in some bacteria, preventing death while experiencing stressful conditions (54–56). The vulnerable strain also upregulates the cytochrome bd complex, an alternative terminal oxidase to cytochrome o ubiquinol oxidase with a high affinity for oxygen that is used in microaerobic conditions (57). It has also been shown to be induced concurrently with growth arrest (58).

Combined with its downregulation of cytochrome o ubiquinol, this suggests the vulnerable strain utilizes alternative, stress-induced respiratory pathways under thermal stress at 36°C. The vulnerable strain’s apparent growth arrest indicates that elevated temperatures put the vulnerable strain under a higher degree of stress than the resistant strain, which instead increases cell proliferation.

The resistant strain exhibits mechanisms to respond to thermal stress without arresting growth. First, regulation of heat shock pathways differed between strains. While both strains upregulated four *hsp20* family heat shock proteins, the resistant strain upregulated them to a significantly higher degree, with an average log2(fold change) of 4.15, compared to the vulnerable strain’s average of 2.43. *Hsp20* proteins are molecular chaperones that bind to unfolded peptides, preventing protein aggregation. Induction levels of these proteins are highly correlated with thermal tolerance . Additional protein chaperones *DnaK*, *DnaJ*, and *GroESL* were only induced in the resistant strain. These proteins bind to unfolded and misfolded proteins, preventing protein aggregation and facilitate refolding . The resistant strain’s more robust induction of heat shock and chaperone proteins likely enhances protein function and stability at elevated temperatures.

#### Cell Membrane Stability

Differential regulation of outer membrane component biosynthesis is also altered by an increase in temperature, as revealed by the KEGG analysis. The upregulation of various nucleotide sugar biosynthesis pathways may contribute to the resistant strain’s higher thermal tolerance. The resistant strain upregulates genes in the pentose phosphate pathway, driving production of ADP-L-glycero-β-D-manno-heptose and CMP-KDO, two main components involved in lipopolysaccharide inner core synthesis (59). Meanwhile, the vulnerable strain downregulates these same genes. Inner core synthesis has been found to be critical to outer membrane thermal stability. *E. coli* mutants with defective inner core synthesis enzymes display greater thermal sensitivity and reduced membrane integrity (60). Induction of inner core synthesis in the resistant strain could reinforce membrane stability at 36°C, while the repression of these same components in the vulnerable strain may contribute to lower membrane stability at 36°C.

While the vulnerable strain represses inner core synthesis, enzymes such as UDP acetylglucosamine epimerase and UDP-N-acetyl-D-galactosamine dehydrogenase, involved in the synthesis of UDP-GalNAc and its derivatives, were upregulated at higher temperatures in the vulnerable strain and downregulated in the resistant strain. UDP-GalNAc initiates O-antigen polymerization in Gram-negative bacteria (61). This may suggest that the vulnerable strain induces O-antigen synthesis, while the resistant strain downregulates these same enzymes, repressing O-antigen synthesis. The resistant strain’s repression of O-antigen synthesis may be due to prioritization of inner core synthesis. LPS inner core and O-antigen synthesis both require undecaprenyl phosphate (Und-P), a lipid phosphate carrier and a limited resource in the cell for which both synthesis processes may compete (62). In *E. coli,* the LPS outer core is truncated under stress and has been shown to block O-antigen ligation, effectively prioritizing inner core synthesis (63). This prioritization may be occurring in the resistant strain, while the vulnerable strain instead continues to focus on O-antigen synthesis.

O-antigen synthesis and variation largely contribute to increased community behaviors, such as quorum sensing and biofilm formation (64,65). Biofilm formation can be induced in Gram-negative bacteria in response to thermal stress, promoting long term survival (66). While biofilm formation assays did not show an increase in biofilm formation at 36°C, the vulnerable strain did have much higher biofilm formation than the resistant strain at both temperatures. A biofilm-based lifestyle in the vulnerable strain may explain why it does not restrict O-antigen synthesis under thermal stress in favor of outer membrane thermal stability, as biofilms’ high heat capacity can dampen the effects of temperature fluctuations (67). A biofilm-based lifestyle also is consistent with a thermal stress response involving growth arrest, as bacteria in biofilms typically have much slower growth rates than when free-living (68). The vulnerable strain upregulates six diguanylate cyclase genes and nine phosphodiesterase genes. Conversely, the resistant strain downregulates eight diguanylate cyclase genes and eighteen phosphodiesterase genes, while only upregulating three phosphodiesterase genes. These enzymes are involved in synthesis and degradation of cyclic-di-GMP, a secondary messenger that induces biofilm formation. In *P. aeruginosa*, diguanylate cyclase deletion mutants are defective in biofilm formation (69,70). Additionally, in the GO enrichment analysis, the vulnerable strain was found to upregulate genes related to viral capsid assembly.

Prophage emergence has been shown to contribute to biofilm formation and rapid evolution of bacterial populations under stressful conditions by contributing to biofilm matrix, cell dispersal, and horizontal gene transfer through phage-induced cell lysis within the biofilm (71).

#### Motility

Cyclic-di-GMP has also been associated with transcriptional changes in flagellar biosynthesis genes (72). The flagellar assembly KEGG pathway had opposite but nonuniform regulation between strains. In the resistant strain, flagellar basal body and hook genes were uniformly and highly upregulated, as was the gene encoding flagellin. Meanwhile, stator protein genes, type III secretion system genes, and regulator protein, *FlgM*, were downregulated. Conversely, the heat-vulnerable strain downregulates basal body, filament, and hook genes and upregulates stator protein genes, the type III secretion system, and *FlgM*. Motility assay results revealed the resistant strain increases motility at 36°C, while the vulnerable strain is nonmotile at both temperatures. However, induction of motility may be unrelated to transcriptional regulation, as multiple bacterial species have shown increased flagellar rotation speed at higher temperatures, without altering the number of flagella (73). Nevertheless, these results may suggest a connection between motility and higher *in vitro* thermal tolerance. For the nonmotile vulnerable strain, transcriptional changes in flagellar assembly genes may instead be related to shifts in community behavior. The vulnerable strain upregulated stator proteins, while the resistant strain downregulated these proteins. In addition to acting as flagellar motors, stator proteins can act as mechanosensors that detect surface contact and initiate biofilm formation (74).

### Comparison with *in vivo* thermal stress response

These same strains were used in an *in vivo* transcriptomic analysis that compared gene expression across temperature regimes when the strains were inside their hosts, with key similarities and differences to our *in vitro* results (29). Both strains downregulated more genes in response to thermal stress *in vivo*, which may suggest that many of the genes upregulated *in vitro* may be unnecessary within the more stable host gut environment. Among the genes still upregulated *in vivo* are heat shock genes and molecular chaperones, which were induced in the resistant strain and not the vulnerable strain. This is similar to what we observed *in vitro*, and may contribute to the resistant strain’s higher thermal tolerance both within and outside of the host. The resistant strain also induced motility genes *in vivo*, consistent with its *in vitro* thermal response. However, unlike our *in vitro* analysis, the vulnerable strain upregulated several motility associated genes *in vivo*, which is in contrast to our current work which revealed a repression of these genes *in vitro*, and a lack of motility at both high and low temperatures.

### Symbiont Stress and Effects on Host Acquisition

As an environmentally acquired symbiont, *Caballeronia’s* survival in the local environment is critical for host acquisition. In legumes, strains of their environmentally acquired microbial symbiont (rhizobia) that have higher growth rates in the local environment are more competitive for nodulation, colonizing host plants more quickly (75).

One experiment found that coinoculation of legumes with different symbiont strains favored a fast-growing, but minimally beneficial symbiont (76). This presents an evolutionary conflict, as less beneficial symbionts may be acquired at higher rates simply due to elevated growth rates under local environmental conditions. As global temperatures increase, *Caballeronia* strains that grow faster at higher temperatures could be acquired more frequently. This mechanism would be further enhanced by interstrain competition. When hosts are provided with a mixed *Caballeronia* symbiont population, strains that are acquired earlier (77), or in larger quantities (78), will dominate the host’s symbiont organ, and in this case, these would be strains that can tolerate higher temperatures. For the strains used in this analysis, higher free-living thermal optimum was correlated with better host outcomes at high temperatures. Thus, in regions exhibiting increasing temperatures, greater acquisition of strains like our resistant strain, V-LZ019, would lead to greater host success. However, some *Caballeronia* strains with higher thermal optima have been found to confer worse host outcomes under thermal stress (19). In these cases, hosts that critically depend on these symbionts for development and survival may suffer. Our analysis reveals that thermal stress responses differ greatly between strains with different thermal optima. Thus, different microbial stress responses may affect host outcomes in the bug-*Caballeronia* symbiosis in the face of climate change, and those symbiont strains with better thermal stress responses, that lead to faster proliferation and cellular stability, may be selected for regardless of host benefits.

This study was subject to some notable limitations. Firstly, the vulnerable strain had an outlier among the 24°C samples (Fig 1). This may contribute to decreased significance levels in the differential expression analysis, likely giving us a less complete picture of the vulnerable strain’s transcriptional response to thermal stress compared to the resistant strain (Fig 2). Furthermore, the symbiont strains’ respective thermal optima were determined in nutrient-rich broth under laboratory conditions. Elucidation of responses to thermal stress in more naturalistic environmental conditions (*i.e.,* when in soil near plants or when in frass excreted by host insects) is an important next step to understanding these bacteria’s environmental performance and potential for shifts in host acquisition under alternative temperature regimes.

## Conclusions

Overall, different thermal response strategies are associated with different thermal tolerances between symbiont strains. The resistant strain’s induction of outer membrane thermal stability, motility, and robust molecular chaperone induction allow it to increase growth at 36°C compared to 24°C through induction of central metabolism, protein synthesis, and cell cycle machinery. Meanwhile, the vulnerable strain’s biofilm-based lifestyle may compromise the thermal stability of its outer membrane. This, combined with weaker induction of molecular chaperones, decreases thermal tolerance, causing growth arrest to promote long term survival. These results may indicate that biofilm-based thermal stress responses in environmentally acquired symbionts may compromise growth and thus host acquisition rates at higher temperatures. Meanwhile, robust motility, chaperone induction, and prioritization of LPS inner core synthesis may be associated with increased growth and host acquisition. As climates warm, these differential thermal stress responses may drive changes in partner-host associations and possible host outcomes in the bug-*Caballeronia* symbiosis.

## Acknowledgments

We would like to thank Derek Wu for help with the experimental design of the motility assay and the rest of the Gerardo and De Roode labs for laboratory help and the use of shared space.

## Author Contributions

PTS conceived the idea and designed the research; PTS collected transcript data; AN conducted and collected data from motility and biofilm assays; AN analyzed data; PTS guided data analysis and verified results; AN wrote the manuscript; AR provided funding for transcriptome sequencing; NMG supervised the project, providing critical guidance to project conceptualization and data analysis; All authors contributed to manuscript revisions.

## Data Availability

Transcript data is available on NCBI SRA under BioProject PRJNA1089548, BioSamples SAMN40544247–SAMN40544264. R code, biofilm and motility assay results, and the Deseq2 output files are available in the supplement.

